# Discrete Ricci curvatures capture age-related changes in human brain functional connectivity networks

**DOI:** 10.1101/2022.12.07.519514

**Authors:** Yasharth Yadav, Pavithra Elumalai, Nitin Williams, Jürgen Jost, Areejit Samal

## Abstract

Geometry-inspired notions of discrete Ricci curvature have been successfully used as markers of disrupted brain connectivity in neuropsychiatric disorders, but their ability to characterize age-related changes in functional connectivity is unexplored. Here, we apply Forman-Ricci curvature and Ollivier-Ricci curvature to compare functional connectivity networks of healthy young and older subjects from the Max Planck Institute Leipzig Study for Mind-Body-Emotion Interactions (MPI-LEMON) dataset (*N* = 225). We found that both Forman-Ricci curvature and Ollivier-Ricci curvature can capture whole-brain and region-level age-related differences in functional connectivity. Meta-analysis decoding demonstrated that those brain regions with age-related curvature differences were associated with cognitive domains known to manifest age-related changes – movement, affective processing and somatosensory processing. Moreover, the curvature values of some brain regions showing age-related differences exhibited correlations with behavioral scores of affective processing. Finally, we found an overlap between brain regions showing age-related curvature differences and those brain regions whose non-invasive stimulation resulted in improved movement performance in older adults. These results suggest that both Forman-Ricci curvature and Ollivier-Ricci curvature correctly identify brain regions that are known to be functionally or clinically relevant. Our results add to a growing body of evidence demonstrating the sensitivity of discrete Ricci curvature measures to changes in the organisation of functional connectivity networks, both in health and disease.

## INTRODUCTION

The proportion of older adults or elderly is increasing across the world. Globally, the number of adults aged 65 years or above surpassed the number of children under the age of 5 for the first time in 2018. Further, the share of the global population aged 65 years or above is projected to rise from 10% in 2022 to 16% in 2050 [1, 2]. This upward shift in the age distribution of the global population makes it crucial [3] to identify the neuronal correlates of age-related decline in cognition, perception and motor performance. The advent of cutting-edge neuroimaging technologies has facilitated the study of anatomical and functional changes in the human brain during the healthy aging process [4, 5]. Specifically, studies employing functional magnetic resonance imaging (fMRI) methods [6–8] have shed light on the altered activity of brain regions that are necessary for optimal cognitive performance in elderly individuals, including the prefrontal, medial temporal and parietal cortices [4, 9, 10]. There is also an increasing interest in the development of novel intervention strategies for reversing the effects of age-related functional decline. Non-invasive brain stimulation techniques such as transcranial direct current stimulation (tDCS), transcranial alternating current stimulation (tACS), and transcranial magnetic stimulation (TMS) offer an attractive option to modulate neuronal plasticity and improve learning processes in older individuals [11–13]. Neuroimaging techniques such as fMRI can not only record the activity of individual regions in the human brain, but also capture the correlations between the activities of these brain regions [14], thereby revealing the underlying functional connectivity networks (FCNs). Numerous efforts have been undertaken to characterize age-related changes in FCNs using concepts and tools from network science and graph theory [15]. Remarkably, graph-theoretic studies on resting-state fMRI (rs-fMRI) datasets of healthy young and healthy older individuals have demonstrated age-related changes in several networkbased measures including clustering coefficient, shortest path length, global efficiency, local efficiency and modularity [16–20]. Recent graph-theoretic studies have combined data from diffusion tensor imaging (DTI) and rs-fMRI scans of healthy aging participants in the Max Planck Institute Leipzig Study for Mind-Body-Emotion Interactions (MPI-LEMON) [21] to analyze the influence of white matter hyperintensities (WMH) and global fractional anisotropy (gFA) on resting-state FCNs [22, 23].

In recent years, several geometric notions of *discrete Ricci curvature* [24–27] have been introduced as novel tools for the analysis of complex networks [28, 29]. Discrete Ricci curvatures allow for a rigorous characterization of network structure by accounting for higher-order correlations in a network [30], and have found important applications such as community detection in networks [31–33] and indicators of critical events in financial markets [34, 35]. Forman–Ricci curvature (FRC) [24, 26] and Ollivier–Ricci curvature (ORC) [25] are the two commonly-used notions of discrete Ricci curvature for analysis of complex real-world networks. Multiple studies have utilizied FRC and ORC to measure changes in structural and functional brain connectivity during neurodevelopmental disorders [36–41]. For example, Chatterjee *et al.* [39] revealed differences in FRC in the FCNs of individuals with attention deficit hyperactivity disorder (ADHD) compared to the FCNs of healthy controls. Recently, Elumalai *et al.* [40] demonstrated that FRC can capture differences in FCNs of individuals with autism spectrum disorder (ASD) compared to FCNs of typically developing controls, and could be used to identify brain regions clinically relevant to ASD. Additionally, Simhal *et al.* [41] used ORC to characterize changes in white matter connectivity after intravenous cord blood infusion in children with ASD. FRC and ORC have also been applied to brain networks of healthy populations. Specifically, Lohmann *et al.* [42] applied FRC to predict the intelligence of healthy human subjects using fMRI data, and Farooq *et al.* [36] applied ORC to identify changes in brain structural connectivity during healthy aging. However, no previous study has assessed the ability of discrete Ricci curvatures to characterize FCN alterations during healthy aging.

In the present work, we apply FRC and ORC to compare resting-state FCNs of healthy young and healthy elderly individuals. We constructed resting-state FCNs by applying a standard fMRI preprocessing pipeline [40] to raw rs-fMRI images of 225 healthy participants in the MPI-LEMON dataset [21]. We estimated whole-brain and region-level FRC and ORC differences between the young and elderly age groups. Next, we used meta-analysis decoding [43, 44] to determine whether discrete Ricci curvatures can identify the cognitive and behavioral domains that are typically associated with healthy aging. Finally, we performed two post-hoc analyses based on the results of the meta-analysis decoding. For the first post-hoc analysis, we determined whether the node curvatures of brain regions with age-related differences were correlated with performance in behavioral tests of affective processing. Affective processing is known to change with age [45]. For the second post-hoc analysis, we compared the set of brain regions showing age-related curvature differences, to the set of brain regions whose non-invasive stimulation with tDCS, tACS or TMS is known to improve motor performance in healthy older adults. Motor performance is also known to change with age [46, 47].

## MATERIALS AND METHODS

We applied Forman-Ricci curvature (FRC) and Ollivier-Ricci curvature (ORC) to measure age-related changes in resting-state functional connectivity networks (FCNs). We constructed the FCNs from raw rs-fMRI images of the subjects acquired from the MPI-LEMON dataset [21]. We used the CONN functional connectivity toolbox [48] to spatially and temporally preprocess the raw rs-fMRI images. Next, we parcellated each preprocessed image into 200 regions of interest (ROIs) or nodes using the Schaefer atlas [49] and generated a 200 × 200 functional connectivity (FC) matrix for each subject. We constructed the FCNs for each subject using a two-step filtering approach, comprising maximum spanning tree (MST) followed by sparsity-based thresholding. Note that the preprocessing pipeline for raw rs-fMRI images and methodology used to construct the FCNs in this work is identical to our previous work on autism spectrum disorder [40]. In the following, we describe in detail, the procedure we employed to construct and analyze the FCNs.

### Participants and imaging dataset

We obtained raw rs-fMRI and anatomical data for 228 participants from the MPI-LEMON dataset [21], comprising 154 young individuals from 20 – 35 years, and 74 elderly individuals from 59 – 77 years. Middle-aged individuals are not included in the MPI-LEMON dataset. We downloaded all the raw rs-fMRI and anatomical data from the ‘Functional Connectomes Project International Neuroimaging Data-Sharing Initiative / Child Mind Institute’ available at the following link: http://fcon_1000.projects.nitrc.org/indi/retro/MPI_LEMON.html. Next, we excluded the subjects with corrupted or missing raw files in the MPI-LEMON dataset, resulting in 225 subjects (young group: 153 subjects, elderly group: 72 subjects) that we included for subsequent analyses. Note that the participants included in the MPI-LEMON dataset provided written informed consent prior to any data acquisition, including agreement to share their data anonymously.

### Raw fMRI data preprocessing

We preprocessed the raw rs-fMRI scans from the MPI-LEMON dataset using the CONN functional connectivity toolbox [48]. The preprocessing pipeline used in this study is identical to our previous work on autism spectrum disorder [40], and we have published earlier a protocol video explaining this pipeline which is available at: https://youtu.be/ch7-dOA-Vlo.

We spatially preprocessed the raw rs-fMRI scans in four major steps: (i) motion correction, (ii) slice-timing correction, (iii) outlier detection, and (iv) structural and functional segmentation and normalization.

For the motion correction step, we co-registered the raw fMRI scans to the first scan of the first session. We corrected the fMRI scans for motion using six rigid body transformation parameters (three translations and three rotations) that are a part of the SPM12 realign and unwarp procedure [50]. For the slice-timing correction step, we corrected temporal misalignment between different slices of the fMRI scans using the SPM12 slice-timing correction procedure [51]. For the outlier detection step, we used Artifact Detection Tools (ART)-based outlier detection to mark acquisitions as outliers if they were found to have framewise displacement greater than 0.5 mm or global BOLD signal changes greater than 3 standard deviations. For the segmentation and normalization step, we normalized the images into the Montreal Neurological Institute (MNI) space using the standard procedure [52]. Subsequently, we segmented the brain into grey matter, white matter and cerebrospinal fluid (CSF) areas using raw T1-weighted volumes of the anatomical scans and mean BOLD signal of the fMRI scans as reference.

With the CONN toolbox, we extracted the BOLD signals (time series) for each voxel after spatially preprocessing the raw rs-fMRI scans, and performed a temporal preprocessing or denoising step to reduce existing motion effects from BOLD signals. First, we applied an anatomical component-based noise correction procedure (aCompCor), which includes a linear regression step. This step resulted in the removal of 5 possible noise components [53] each from white matter and CSF areas, 12 possible noise components due to estimated subject motion parameters and their first-order derivatives [54], and 1 noise component from outlier scans detected earlier (scrubbing) [55]. Second, we removed temporal frequencies less than 0.008 Hz from the BOLD signals using a high-pass filtering approach.

All the 225 subjects passed the quality assessment checks provided by the CONN toolbox [56] (see **Supplementary Table S1**). Finally, we used the preprocessed fMRI scans of these 225 subjects to construct FC matrices and perform network analysis.

### ROI definition and ROI time series estimation

After spatially and temporally preprocessing the raw rs-fMRI scans of the 225 subjects, we partitioned the voxels into cortical parcels so as to define nodes or ROIs with interpretable neurobiological meaning [57] and reduce the computational load of further analysis. For this purpose, we used a cortical parcellation atlas provided by Schaefer *et al.* [49], which parcellates the brain into 200 distinct ROIs or nodes. Apart from assigning each voxel to one of 200 ROIs, the Schaefer atlas also provides a mapping between each ROI and one of seven resting state networks (RSNs), namely, ‘default network’, ‘somatomotor network’, ‘dorsal attention network’, ‘salience/ventral attention network’, ‘visual network’, ‘limbic network’ and ‘control network’. With the CONN toolbox, we determined the time series corresponding to each ROI by computing the average time series of its constituent voxels.

### Functional connectivity network construction

We constructed a 200 × 200 FC matrix for each of the 225 subjects, by computing the Pearson correlation coefficient between all pairs of ROI time series. The resulting FC matrix can be considered a complete, weighted and undirected network with 200 nodes, with the weight of each edge determined by the Pearson correlation coefficient between the time series of its corresponding pair of nodes. We used the FC matrix of each of the 225 subjects to construct the FCNs, using a two-step procedure.

First, we determined the maximum spanning tree (MST) of the FC matrix using Kruskal’s algorithm [58]. The MST for a weighted graph with *n* nodes is an acyclic graph with (*n* – 1) edges which includes all the nodes of the graph. Using an MST-based filtering approach ensures that the resulting FCN is a connected graph and it captures the highly correlated nodes from the FC matrix.

Second, we used a sparsity-based thresholding procedure, wherein edges from the FC matrix were iteratively added to the MST in decreasing order of their correlation values until we generated a network with the desired edge density. We binarized the thresholded network by removing the edge weights in order to obtain an unweighted, undirected and connected FCN with desired edge density [16, 59]. Using a sparsity-based filtering approach ensures that the resulting FCNs for different subjects have the same number of edges, which enables a direct mathematical comparison of discrete Ricci curvatures and other network properties across subjects [59–61]. We remark that such a two-step procedure with MST followed by sparsity-based thresholding was used previously by Achard *et al.* [62] and Elumalai *et al.* [40] to construct resting-state FCNs from FC matrices.

In this work, we computed discrete Ricci curvatures and other network properties over the range of edge densities 0.02 – 0.5 or 2% – 50% edges, with an increment of 0.01 or 1% edges [59, 63, 64]. Therefore, we constructed 49 FCNs for each of the 225 subjects from the MPI-LEMON dataset. We have made all the 200 × 200 FCNs for 225 subjects across 49 graph densities or thresholds publically available via a GitHub repository: https://github.com/asamallab/Curvature-FCN-Aging.

### Discrete Ricci curvatures

Each FCN generated in our work can be represented as an unweighted and undirected graph or network *G* = (*V, E*), where *V* is the set of vertices or nodes in G and E is the set of edges or links in G. Classically, Ricci curvature is defined on tangent vectors on smooth manifolds [65], whereas for discrete objects such as networks, Ricci curvature is naturally defined on the edges. While both FRC [24, 26] and ORC [25] are discrete analogues of the classical Ricci curvature, the definitions for FRC and ORC are non-equivalent, and therefore, the two discrete versions capture distinct properties of the classical Ricci curvature. Notably, FRC captures the geodesic dispersal property of the classical Ricci curvature whereas ORC captures the volume growth property of the classical Ricci curvature [27]. Below, we present definitions of the two notions of discrete Ricci curvature, specific to an unweighted and undirected network, employed in our work.

#### Forman-Ricci curvature

Forman [24] introduced a discretization of Ricci curvature for a particular class of geometric objects known as *CW complexes*, which are widely studied within the field of algebraic topology. Some of us adapted the notion of Forman-Ricci curvature (FRC) to undirected graphs [26], which are equivalent to 1-dimensional CW complexes. However, a remarkable property of Forman’s discretization is that it can be defined for CW complexes in any arbitrary number of dimensions, which allows for a rigorous means to account for higher-order interactions in complex systems. Notably, we utilized this property in a subsequent work [27] and defined a modified version of FRC, also known as *augmented* FRC. Formally, for a graph G where weights are assigned to vertices, edges and triangular faces (that is, cycles of length 3), the augmented FRC of an edge *e* is defined as

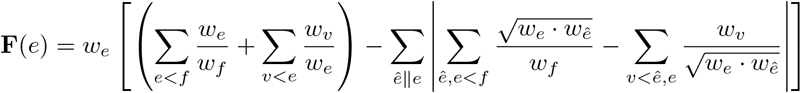

where *w_e_* is the weight of edge *e, w_v_* is the weight of vertex *v, w_f_* is the weight of triangular face *f. σ* < *τ* means that the *σ* is a lower dimensional face of *τ*, || means that two cells are parallel, i.e. they either share a higher dimensional face or a lower dimensional face, but not both. Augmented FRC accounts for cycles of length 3 in a graph while neglecting cycles of length 4 or higher.

If the graph *G* is unweighted, then all the vertices, edges and triangular faces are assigned weight equal to 1, and the augmented FRC reduces to the following simple combinatorial expression

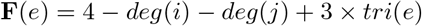

where *deg*(*i*) and *deg*(*j*) are the degrees of nodes *i* and *j*, respectively, that are anchored to the edge e and *tri(e)* is the number of triangular faces that contain the edge *e*. We refer to augmented FRC simply as FRC throughout the main text. Intuitively, FRC quantifies the extent of the spread of information around the ends of an edge in a network. A more negative value of FRC of an edge indicates a higher amount of information spread around its ends.

#### Ollivier-Ricci curvature

For an edge *e* between nodes *i* and *j* in an unweighted and undirected graph *G*, ORC is measured by comparing the minimal cost of transporting a mass distribution over the neighbors of *i* and *j* with the distance between *i* and *j* itself [25, 66]. Formally,

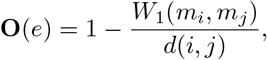

where *m_i_* and *m_j_* are the discrete probability measures defined on nodes *i* and *j*, respectively, and *d*(*i,j*) is the distance between *i* and *j*. Since *G* is an unweighted graph, *d*(*i,j*) is equal to the number of edges contained in the shortest path connecting *i* and *j*. *W*_1_ is the trasportation distance between *m_i_* and *m_j_*, also known as the Wasserstein distance [67], which is defined as

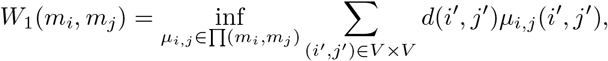

where ∏(*m_i_, m_j_*) is the set of probability measures *μ_i,j_* that satisfy

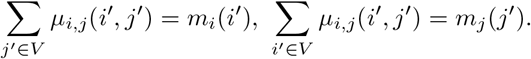

The above equation gives all the possibile transportations of measure *m_i_* to *m_j_*. The Wasserstein distance *W*_1_(*m_i_, m_j_*) is the minimal cost of transporting *m_i_* to *m_j_*. The probability distribution *m_i_* is considered to be uniform over the neighboring nodes of *i* [68]. Note that the computation of ORC can be formulated as a linear programming problem [66].

### Global and node-level network analyses

For each of the 225 subjects in the MPI-LEMON dataset included in this study, we generated 49 unweighted and undirected FCNs with varying edge densities from 2% – 50% with an increment of 1%. Subsequently, we compared the global or whole-brain characteristics of the FCNs, at each edge density, across the two groups (young and elderly) using FRC and ORC. Specifically, we computed the average FRC of edges and average ORC of edges, for each of the 49 networks, for each of the 225 subjects included in this study. Additionally, we computed eight other global network measures namely, clique number, average clustering coefficient, global efficiency [69], average node betweenness centrality, average local efficiency [69], average shortest path length, modularity [70], and assortativity. We provide the definitions of theses global network measures used to characterize the FCNs in the Supplementary Information.

Next, we compared the node or region-level characteristics of the FCNs, at each edge density, across the young and elderly groups. Specifically, we computed the node FRC and node ORC for the 200 nodes, for each of the 49 networks, for each of the 225 subjects included in this study. The notion of node Ricci curvature is analogous to the notion of scalar curvature in Riemannian geometry [65] and is computed as the sum of the Ricci curvatures for the edges incident on a given node [27].

The computer codes used to compute the global and node-level network characteristics including FRC and ORC are publicly available via a GitHub repository: https://github.com/asamallab/Curvature-FCN-Aging. The standard network measures were computed using the Python package NetworkX [71].

### Neurosynth meta-analysis decoding

In order to determine the cognitive and behavioral relevance of the results from node-level comparisons across the young and elderly groups, we used Neurosynth meta-analysis decoding [43, 44]. First, we used the Neurosynth metaanalysis tool to find terms related to cognition, perception and behavior corresponding to the centroid coordinates of each of the 200 Schaefer ROIs. Second, we assigned each of the nodes or ROIs with significant between-group differences in FRC or ORC to their respective RSN, as defined by the Schaefer atlas. The ROIs were assigned to 1 of 7 such RSNs, since the Schaefer atlas is divided into 7 RSNs. Third, for a given RSN, we computed the number of occurrences of the terms associated with the significant ROIs within that RSN. Fourth, we determined the statistical significance of the number of occurrences of the different terms associated with each RSN. Note that the above steps were performed separately for each RSN.

### Statistical analyses

We employed a two-tailed two-sample t-test to estimate the differences in average edge curvatures and other global network measures between the young and elderly groups throughout the 49 FCN densities in the range 2 – 50% considered in this work. In order to estimate the between-group differences in node curvatures, we first computed the area under the curve (AUC) for the FRC or ORC of a given node across the 49 graph densities [16, 63]. Subsequently, we employed a two-tailed two-sample *t*-test on the AUCs of the discrete curvatures for each of the 200 nodes of the FCNs.

In order to determine the statistical significance of the number of occurrences of each term in the Neurosynth metaanalysis decoding, we first computed the number of occurrences of the same terms corresponding to equal number of randomly selected nodes (surrogate ROIs). For example, if we identified 66 nodes with significant between-group differences in FRC, then we randomly selected a set of 66 nodes out of the 200 nodes in the Schaefer atlas. We obtained a null distribution for the number of occurrences of each term by generating 1000 sets of randomly selected nodes. Subsequently, we calculated the *z*-score corresponding to the number of occurrences of each term associated with the subset of original ROIs. Finally, we converted the *z*-scores to *p*-values, assuming a normal distribution [40, 44].

In order to correct for multiple comparisons and control the occurrence of false positives, we employed a false discovery rate (FDR) correction [72] to adjust the *p*-values for each of the statistical tests described above. We set the threshold *α* for these FDR corrections to 0.05. These statistical tests were performed using Python packages SciPy [73] and statsmodels [74].

### Correlation of node curvatures with cognitive and behavioral scores

Neurosynth meta-analysis decoding revealed cognitive and behavioral domains associated with those brain regions whose node curvatures were different between young and elderly participants. We performed a correlation analysis to determine the extent to which curvatures of individual brain regions could account for variation in performance on tests measuring ability in cognitive or behavioral domains linked to those regions. Specifically, if the Neurosynth metaanalysis for a given RSN revealed a particular cognitive or behavioral domain, then we acquired all the phenotypic test scores associated with that domain for each of the 225 subjects in the MPI-LEMON dataset. Next, we computed Spearman’s correlation between the node FRC (or node ORC) of all the significant regions in that RSN and the phenotypic scores provided in each of the cognitive or behavioral tests. We used Spearman rather than Pearson correlation since some of the test scores are expressed on an ordinal rather than continuous scale. Finally, we performed FDR correction to adjust the *p*-values of the estimated correlations. FDR correction was applied to *p*-values of each cognitive or behavioral test, separately. Note that if a particular cognitive or behavioral test had multiple phenotypic scores, then the FDR correction was applied across all the phenotypic scores within that test.

### Literature search of non-invasive brain stimulation studies in healthy elderly individuals

We performed a literature search on PubMed to identify scientific papers reporting the effect of three non-invasive brain stimulation techniques namely, tDCS, tACS and TMS on motor performance in healthy elderly individuals. Then, we assessed the results reported in these papers to identify brain regions that show evidence for improvements in motor performance of elderly individuals after non-invasive stimulation. The following search query was used in PubMed: ((transcranial magnetic stimulation) OR (TMS) OR (transcranial direct current stimulation) OR (tDCS) OR (transcranial alternating current stimulation) OR (tACS)) AND ((healthy old) OR (healthy elderly) OR (healthy aging) OR (healthy ageing)) AND ((motor function) OR (motor performance) OR (motor skill) OR (movement)). This PubMed search was performed in October 2022, and returned a list of 1659 articles.

We followed a three-stage procedure to identify relevant articles from the list of 1659 articles returned by the PubMed search (see **Figure 1)**. First, we checked all the meta-analysis papers studying the effects of tDCS, tACS or TMS on motor performance in elderly individuals [76–79] to identify articles that were missing from the list of 1659 articles. Second, we filtered the relevant articles from the list of 1659 articles based on title and abstract. Third, we classified the articles according to the non-invasive stimulation technique employed (tDCS, tACS or TMS) and checked the full text of the articles for relevance. We used the following inclusion criteria to judge article relevance: (1) studies on healthy elderly individuals, (2) studies that employ non-invasive brain stimulation, namely tDCS, tACS and TMS, (3) studies that analyze the effect of non-invasive stimulation techniques on motor performance, as measured during movement-related tasks such as finger tapping tasks (FTT), sequence learning tasks and serial reaction time tasks (SRTT) and their variants, and (4) studies that are peer-reviewed. We used the following exclusion criteria to judge article relevance: (1) review articles, (2) articles presented in languages other than English, (3) studies that did not employ non-invasive stimulation, (4) studies that investigate new protocols for brain stimulation (5) studies on diseased populations, including patients with Alzheimer’s Disease, Parkinson’s Disease or Stroke, (6) studies on non-human species, e.g., rats, (7) studies that employed non-invasive stimulation on children or young adults, (8) studies that analyze the effect of non-invasive stimulation on domains other than motor performance, and (9) studies that analyze the effect of non-invasive stimulation on neuronal activity, neuroplasticity or neurophysiological processes. A given article was considered as relevant only if all the inclusion criteria were satisfied, whereas it was considered as not relevant if at least one exclusion criterion was satisfied.

**FIG. 1.**
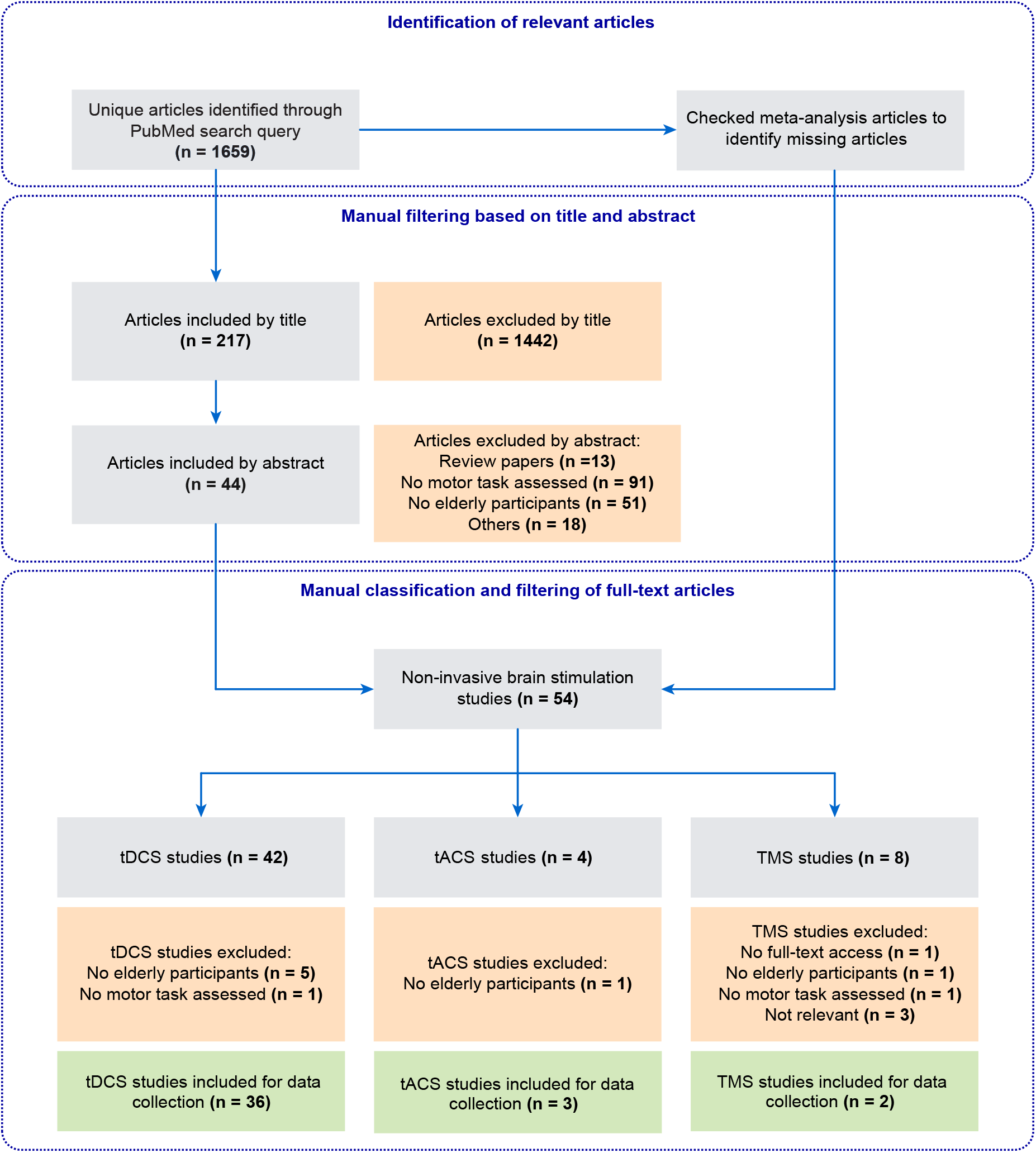
Summary of the workflow used to collect and classify data from non-invasive brain stimulation experiments. The workflow is presented according to PRISMA statement [75]. We obtained a list of 1659 articles from PubMed that report the effects of non-invasive brain stimulation on motor performance during healthy aging. We followed a three-stage procedure to extract relevant articles from the PubMed search. First, we identified missing articles from the original PubMed search by checking meta-analysis papers studying the effects of non-invasive stimulation on motor performance during healthy aging. Second, we filtered the articles based on title and abstract. Third, we classified the articles according to the non-invasive stimulation technique (tDCS/tACS/TMS) and checked the full-text of these articles for relevant data. Finally, we extracted experimental data from 36 tDCS studies, 3 tACS studies and 2 TMS studies.

The above procedure resulted in 36 eligible articles that employed tDCS, 3 eligible articles that employed tACS, and 2 eligible articles that employed TMS in their experiments. We extracted the following data from the eligible articles: PMID, author, publication year, participant information (which includes number of participants and mean age of the participants), type of stimulation (for example, anodal tDCS, cathodal tDCS, bilateral tDCS, rTMS), target brain region, stimulation parameters (such as stimulation intensity for tDCS and tACS, pulses per session and inter-train interval for TMS, number of sessions, and treatment duration), outcome measures used to assess motor performance after stimulation, results of stimulation and adverse effects of stimulation on experimental group or control group (if applicable). **Supplementary Table S2** provides the data extracted from the tDCS, tACS and TMS studies respectively. From the above data, we combined information about the target region, outcome measures and results of tDCS/tACS/TMS stimulation as measured using the outcome measures in each study to identify the set of target regions whose non-invasive stimulation resulted in improved motor performance of healthy older individuals. These target regions were used in a subsequent analysis to determine whether FRC or ORC can identify brain regions whose non-invasive stimulation shows evidence for improvements in motor performance of healthy older individuals.

### Determining overlap between brain regions with age-related differences in curvature and target regions in non-invasive stimulation studies

The target regions in our collected list of non-invasive brain stimulation experiments correspond to the ROIs in the Brodmann atlas [80], whereas the brain regions with age-related differences in FRC or ORC correspond to the ROIs in the Schaefer 200 atlas. Hence, we used the following procedure to enable a systematic comparison of the age-related differences in discrete Ricci curvatures with results from previous non-invasive stimulation studies. First, we mapped each Schaefer ROI to a corresponding Brodmann ROI. The mapping was performed using the MRIcron tool [81] by identifying the Brodmann area that contains the MNI centroid coordinates of a given Schaefer ROI. Note that a similar mapping approach was employed in our previous study by Elumalai *et al.* [40]. Second, we determined the set of Brodmann ROIs corresponding to the target regions in non-invasive brain stimulation experiments that show evidence for improvements in motor performance in healthy elderly. Third, we identified the set of Schaefer ROIs that were mapped to these Brodmann ROIs. Fourth, we identified the Schaefer ROIs within this set that also showed significant age-related differences in FRC or ORC.

## RESULTS

In this work, we employed two geometry-inspired graph measures namely, FRC and ORC, to investigate the wholebrain and region-level differences in resting state FCNs during healthy aging. We preprocessed raw rs-fMRI images of 153 healthy, young individuals in the age range 20 – 35 years and 72 healthy, elderly individuals in the age range 59 – 77 years from the MPI-LEMON dataset. Next, we used the Schaefer atlas [49] to parcellate the preprocessed rs-fMRI scan of each subject into 200 ROIs, and calculated the Pearson correlation coefficient between all pairs of ROI time series, yielding a 200 × 200 FC matrix. Finally, we generated 49 FCNs for each subject across the range of edge densities 0.02 – 0.5 or 2% – 50% edges, with an increment of 0.01 or 1% edges using a two-step edge filtering approach comprising MST and sparsity-based thresholding. The detailed methodology to construct and analyze the FCNs is described in **Materials and Methods**.

### Whole-brain differences in functional connectivity networks

We computed the average edge FRC and average edge ORC for the 49 FCNs across edge densities 2% – 50% for each of the 225 subjects to evaluate the age-related brain-wide changes in FCNs between the healthy young and healthy elderly groups. Subsequently, we applied a two-tailed two-sample t-test with FDR correction (see **Materials and Methods**) to estimate the differences in average edge curvatures between the young and elderly groups at each value of edge density. **Figures 2A** and **2B** show the differences in average edge FRC and average edge ORC, respectively, between the young and elderly individuals over the range of edge densities 2% – 50%. We found that the elderly group had significantly higher values of average edge FRC (*p* < 0.05, FDR-corrected) over the range of edge densities 3% – 50% (**Figure 2A**). Similarly, we found that the elderly group had significantly higher values of average edge ORC over the entire range of graph densities 2% – 50% considered in this study (**Figure 2B**).

**FIG. 2.**
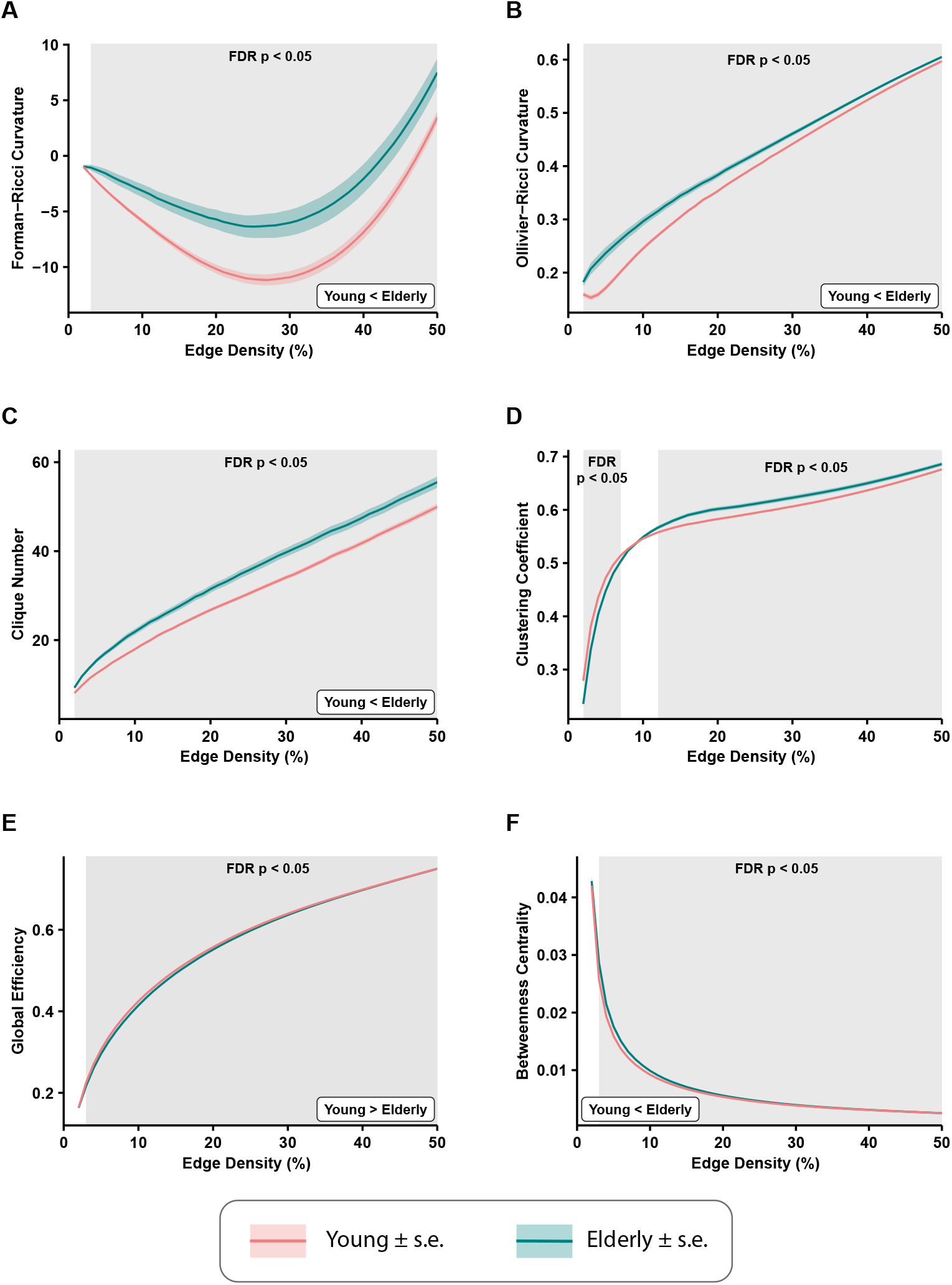
Differences in average edge curvatures and standard global network measures across the functional connectivity networks (FCNs) of 153 young individuals and 72 elderly individuals from the MPI-LEMON dataset. The differences are reported for FCNs over the range of edge densities 0.02 (i.e., 2% edges) and 0.5 (i.e., 50% edges), with an increment of 0.01 (i.e., 1% edges). The shaded regions in each plot correspond to the edge densities where the between-group differences are statistically significant (*p* < 0.05, FDR-corrected). **(A)** Average Forman–Ricci curvature (FRC) of edges is higher in elderly individuals over edge densities 3 – 50%. **(B)** Average Ollivier–Ricci curvature (ORC) of edges is higher in elderly individuals over the entire range of edge densities considered (2 – 50%). **(C)** Clique number is higher in elderly individuals over all edge densities 2 – 50%. **(D)** Average clustering coefficient is higher in young individuals over edge densities 2 – 7%, and higher in elderly individuals over edge densities 12 – 50%. **(E)** Global efficiency is reduced in elderly individuals over edge densities 3 – 50%. **(F)** Average node betweenness centrality is higher in elderly individuals over edge densities 3 – 49%.

We further compared eight other global network measures to evaluate brain-wide changes in FCNs of young and elderly individuals. The eight global network measures include clique number, average clustering coefficient, global efficiency, average node betweenness centrality, average local efficiency, shortest path length, modularity and assortativity. Out of the eight measures, clique number, node betweenness centrality and assortativity have not been applied in previous network-based analyses of rs-fMRI scans in healthy aging. First, we found that the clique number is significantly higher in elderly group (*p* < 0.05, FDR-corrected) over all edge densities considered in this study (**Figure 2C**). Second, we found average clustering coefficient is significantly higher in the young group at edge densities 2% – 7%, whereas it is significantly higher in the elderly group at edge densities 12% – 50% (Figure **2D**). This change in the directionality of the differences in clustering coefficient with increasing edge densities suggests that highly correlated nodes or ROIs tend to cluster in young individuals whereas weakly correlated nodes tend to cluster in elderly individuals. Notably, the group differences in clustering coefficient observed in the present work are consistent with previous studies that have used clustering coefficient to compare the global changes in resting state FCNs in healthy aging cohorts [17, 18]. Third, we found that global efficiency of the FCNs is significantly reduced in elderly individuals over the edge density range 3% – 50% (**Figure 2E**), and our results are consistent with previous graph-theoretic rs-fMRI studies [16–20] in healthy aging populations. Fourth, we found that average node betweenness centrality of the FCNs is significantly higher in elderly individuals over the edge density range 3% – 49% (**Figure 2F**).

Fifth, we found that average local efficiency is significantly higher in the FCNs of the young group compared to the elderly group over the edge density range 1% – 12%, and significantly higher in the elderly group over the edge density range 19% – 49% (**Supplementary Figure S1**). Notably, the differences in average local efficiency observed in the present work are consistent with previous graph-theoretic rs-fMRI studies [16, 19, 20] in healthy aging populations. Sixth, we found that average shortest path length is significantly higher in the elderly group over the edge density range 3% – 50% (**Supplementary Figure S1**), and these differences are consistent with previous graph-theoretic rs-fMRI studies [17] in healthy aging populations. Seventh, we found that modularity is significantly reduced in elderly individuals over the range of edge densities 2% – 9% (**Supplementary Figure S1**), and these differences are consistent with results from previous graph-theoretic rs-fMRI studies [19, 20] in healthy aging populations. Eighth, we found that assortativity is reduced in the elderly group, but the differences are not significant (as suggested by *p* > 0.05, FDR-corrected).

**Supplementary Table S3** lists the FDR-corrected *p*-values corresponding to the group-wise comparisons of discrete Ricci curvatures and standard network measures across all edge densities 2% – 50% considered in this study.

### Region-level differences in functional connectivity networks

As reported in the previous subsection, we found that both FRC and ORC show significant differences between the resting state FCNs of young and elderly individuals at the whole-brain level. Subsequently, we determined how the global differences in discrete Ricci curvatures are distributed across the 200 nodes or ROIs in the brain as defined by the Schaefer atlas (see **Materials and Methods**). First, for each of the 225 subjects, we computed node FRC and node ORC for the 200 nodes in the 49 FCNs across the range of edge densities 2% – 50%. Second, in order to determine the nodes with significant between-group differences in node Ricci curvatures in the FCNs of young and elderly individuals, we computed the Area Under Curve (AUC) of the node curvature values across the 49 edge densities for each of the 200 nodes, and compared these AUCs for each node using a two-tailed two-sample t-test with FDR correction.

**Figures 3A** and **3B** illustrate the nodes or ROIs that display significant between-group differences (*p* < 0.05, FDR-corrected) in FRC and ORC, respectively, of the FCNs of young and elderly individuals. We identified 66 ROIs with significant differences in FRC and 53 ROIs with significant differences in ORC across the young and elderly groups. We found 42 ROIs to be identified by both FRC and ORC. We found that the significant ROIs identified by both FRC and ORC are spread across the 7 RSNs. However, the ROIs with significant differences in node FRC are concentrated in 3 RSNs namely, somatomotor network (33 ROIs), salience/ventral attention network (10 ROIs) and dorsal attention network (10 ROIs). The ROIs with significant differences in node ORC are concentrated in 2 RSNs namely, somatomotor network (27 ROIs) and dorsal attention network (9 ROIs). **Supplementary Table S4** lists the ROIs with significant between-group differences as identified by FRC and ORC, partitioned across the 7 RSNs.

**FIG. 3.**
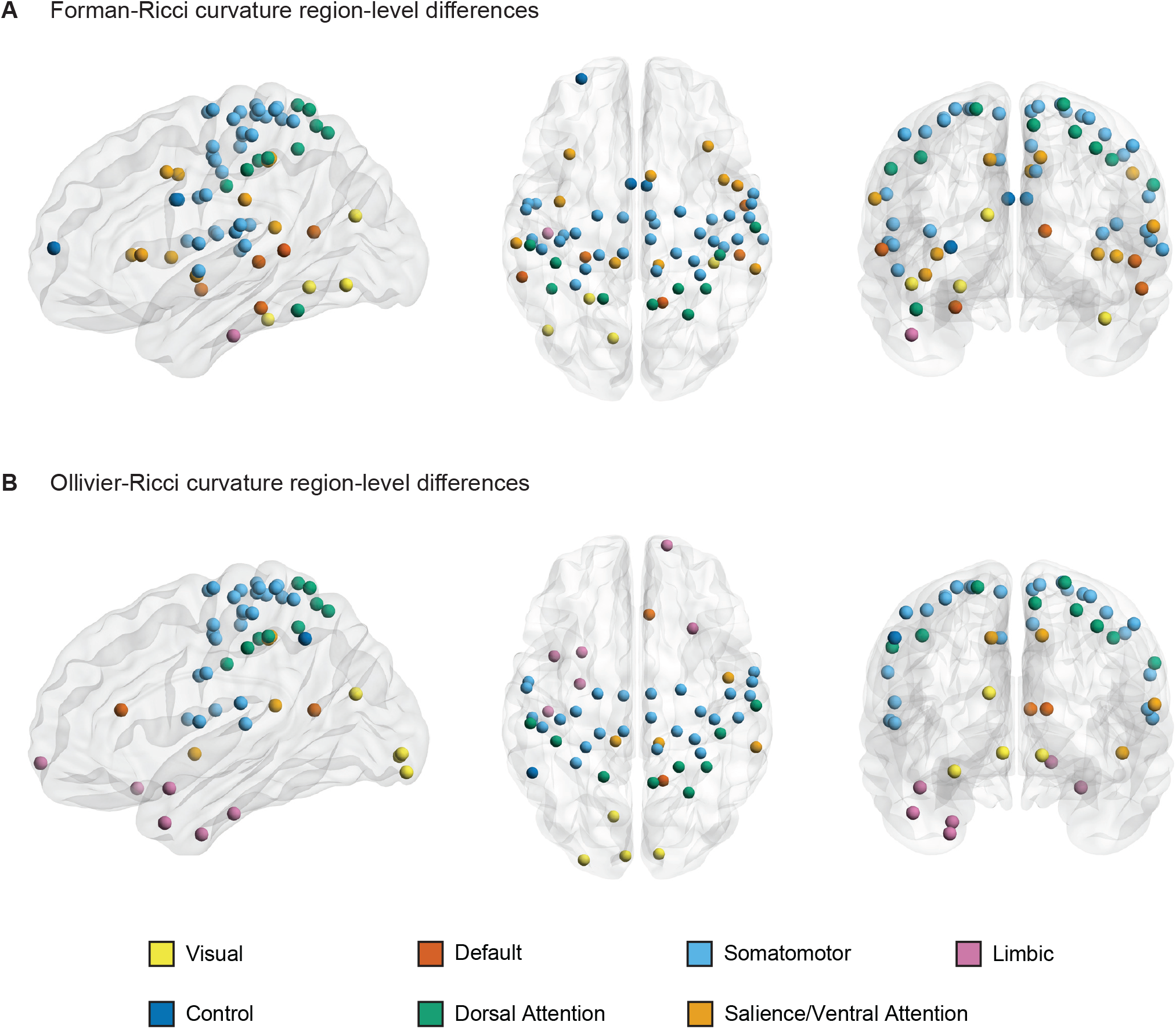
A visual representation of nodes or regions in the brain that show significant differences in discrete Ricci curvatures between the functional connectivity networks (FCNs) of young and elderly individuals (*p* < 0.05, FDR corrected). **(A)** 66 regions that show significant differences in Forman–Ricci curvature (FRC) of the nodes in the FCNs of the young and elderly groups. All the 66 regions display higher values of node FRC for elderly individuals compared to young individuals. **(B)** 53 regions that show significant differences in Ollivier-Ricci curvature (ORC) of the nodes in the FCNs of the young and elderly groups. 41 out of these 53 regions display higher values of node ORC in elderly individuals and the remaining 12 ROIs display higher values of node ORC in young individuals. The nodes are specified by the Schaefer atlas, and color-coded as per the 7 resting state networks (RSNs) listed in the figure legend. This figure was created using BrainNet Viewer [82]

Further, we determined the directionality of the significant differences in node FRC and node ORC between the young and elderly individuals across each of the 200 nodes or ROIs. Recall that at the whole-brain level, both FRC and ORC displayed higher values in elderly individuals compared to young individuals. **Supplementary Table S5** lists the AUCs of the node FRC and node ORC for each of the 200 nodes averaged across the young and elderly groups. We found that all the 66 ROIs with significant between-group differences in node FRC display higher values of node FRC for elderly individuals compared to young individuals. Moreover, we found that 41 out of the 53 ROIs with significant between-group differences in node ORC display higher values of node ORC in elderly individuals and the remaining 12 ROIs display higher values of node ORC in young individuals.

### Behavioral and cognitive relevance of region-level differences

As reported in the previous subsection, we identified 66 nodes or ROIs that show significant between-group differences in FRC (**Figure 3A**) and 53 ROIs that show significant between-group differences in ORC (**Figure 3B**) across the young and elderly groups. We partitioned the set of significant ROIs into 7 subsets based on their respective RSNs as defined by the Schaefer atlas. Subsequently, we determined the cognitive domains associated with the significant ROIs in each RSN using Neurosynth meta-analysis (see **Materials and Methods**). For nodes identified by FRC, we limited the Neurosynth analysis to somatomotor network, salience/ventral attention network and dorsal attention network. For nodes identified by ORC, we limited the Neurosynth analysis to somatomotor network and dorsal attention network. We chose the above-mentioned RSNs since the significant ROIs identified by FRC or ORC are mainly concentrated within these RSNs. Moreover, the sets of significant ROIs in these RSNs are nearly bilaterally symmetrical.

**Figure 4A** shows the word clouds highlighting the behavioral relevance of the significant brain regions associated with three main RSNs identified by FRC, and **Figure 4B** shows the word clouds for the significant brain regions associated with two main RSNs identified by ORC. **Supplementary Table S6** lists the terms associated with the significant regions identified by FRC and ORC across all 7 RSNs. For the nodes identified by FRC, the word cloud for somatomotor network shows terms associated with movement. For salience/ventral attention network, we find terms associated with affective and somatosensory processing. For dorsal attention network, we find terms associated with movement. For the nodes identified by ORC, the word clouds for both somatomotor network and dorsal attention network show terms associated with movement.

**FIG. 4.**
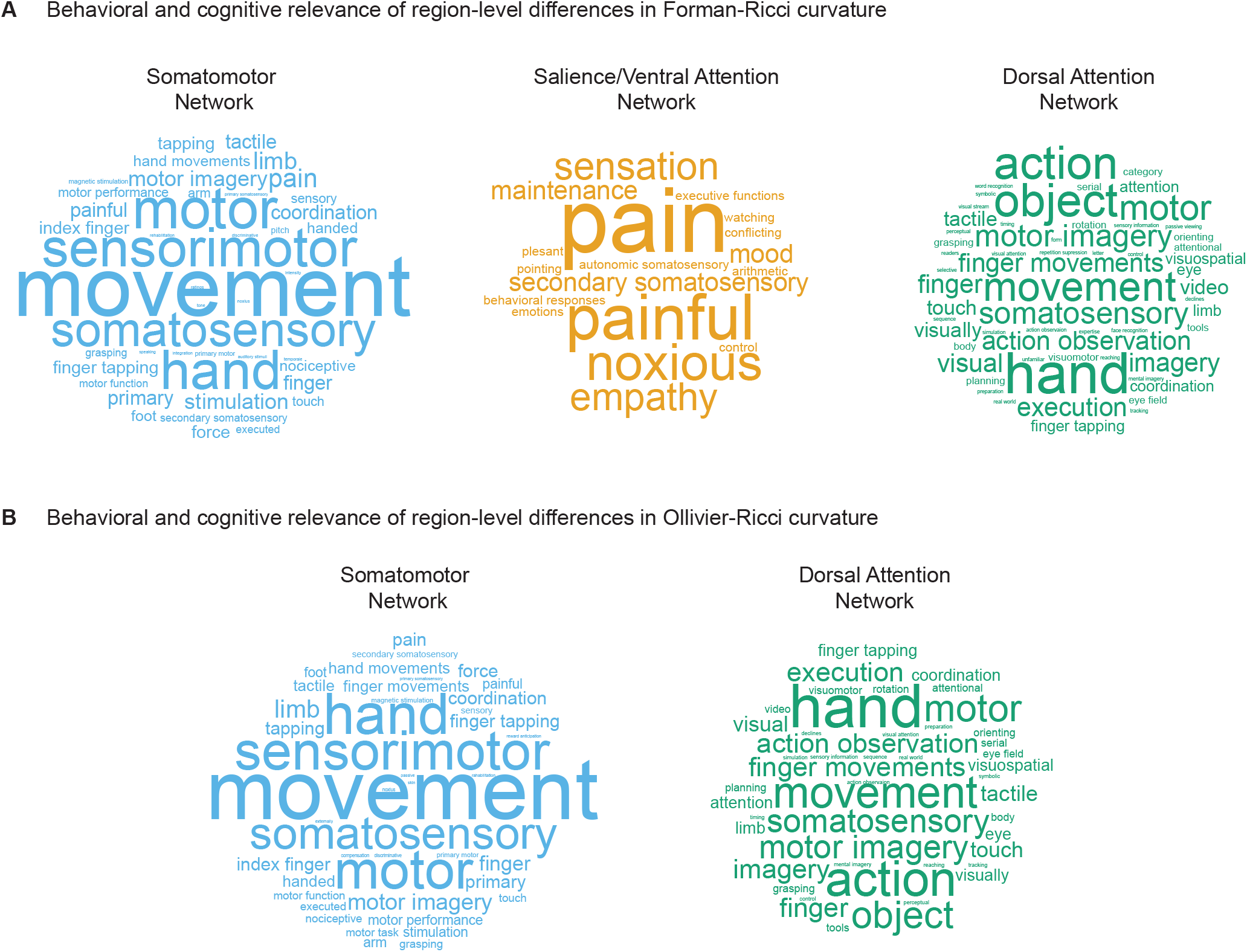
Behavioral and cognitive relevance of age-related region-level differences in curvatures. **(A)** Word clouds displaying cognitive and behavioral terms associated with brain regions having age-related differences in values of Forman-Ricci curvature (FRC), in three RSNs, namely somatomotor network, salience/ventral attention network and dorsal attention network. **(B)** Word clouds displaying cognitive and behavioral terms associated with brain regions having age-related differences in values of Ollivier-Ricci curvature (ORC), in two RSNs, namely somatomotor network and dorsal attention network. The size of the terms in each word cloud is proportional to their frequency of occurrence. Note that size of the terms in each word cloud are scaled separately, and thus, frequencies of occurrence cannot be compared across word clouds. The word clouds in this figure are generated using https://www.wordclouds.com.

### Correlation between curvatures and affective processing-related phenotypic test scores

The Neurosynth meta-analysis revealed that brain regions showing age-related FRC and ORC differences in somatomotor and dorsal attention networks are associated with movement, and brain regions showing age-related FRC differences in salience/ventral attention network are associated with affective and somatosensory processing (**Figure 4)**. To do this, we performed a post-hoc correlation analysis to determine the extent to which node FRC and node ORC of the brain regions with age-related differences in a given RSN, could account for variation in phenotypic test scores corresponding to the cognitive or behavioral domains identified by the Neurosynth meta-analysis for that RSN.

We identified 8 tests related to affective processing, for each of the 225 subjects in the MPI-LEMON dataset, but did not find any tests related to movement or somatosensory processing. In the following, we list these tests related to affective processing: (i) Cognitive Emotion Regulation Questionnaire (CERQ), (ii) Coping Orientations to Problems Experienced (COPE), (iii) Emotion Regulation Questionnaire (ERQ), (iv) Measure of Affect Regulation Style (MARS), (v) Perceived Stress Questionnaire (PSQ), (vi) State-Trait-Angstinventar (STAI-G-X2), (vii) Trait Emotional Intelligence Questionnaire - Short Form (TEIQue-SF), and (vii) Trierer Inventar zum Chronischen Stress (TICS).

We found that node FRC showed significant correlation (p < 0.05, FDR-corrected) only with the TICS phenotypic scores. TICS contains 9 scores that are designed to measure the level of chronic stress that an individual experiences across 9 subdomains including work overload, social overload, pressure to perform, work discontent, excessive demands from work, lack of social recognition, social tension, social isolation, and chronic worrying [83, 84]. Further, TICS contains a separate score called Short Screening Scale for Chronic Stress (SSCS) which was designed as brief chronic stress instrument for applied research and practitioners. Hence, 10 scores are recorded in the TICS test.

Since 10 brain regions in ventral attention network revealed node FRC age-related differences, we performed 10 × 10 = 100 FRC - TICS correlations, out of which 17 correlations were statistically significant (p < 0.05, FDR-corrected). The 17 statistically significant correlations were spread across 7 brain regions, with node FRC of RH_SalVentAttn_PrC_1 being correlated to as many as 6 TICS scores (**Figure 5)**, while the other statistically significant correlations were distributed between RH_SalVentAttn_Med_2, RH_SalVentAttn_Med_1, RH_SalVentAttn_FrOperlns_4, RH_SalVentAttn_TempOccPar_2, LH_SalVentAttn_Med_2 and LH_SalVentAttn_FrOperIns_3 (**Supplementary Figure S2**). All the 17 statistically significant FRC - TICS correlations were negative, ranging between –0.22 and –0.18. Hence, we find that higher node FRC values are associated with lower levels of chronic stress.

**FIG. 5.**
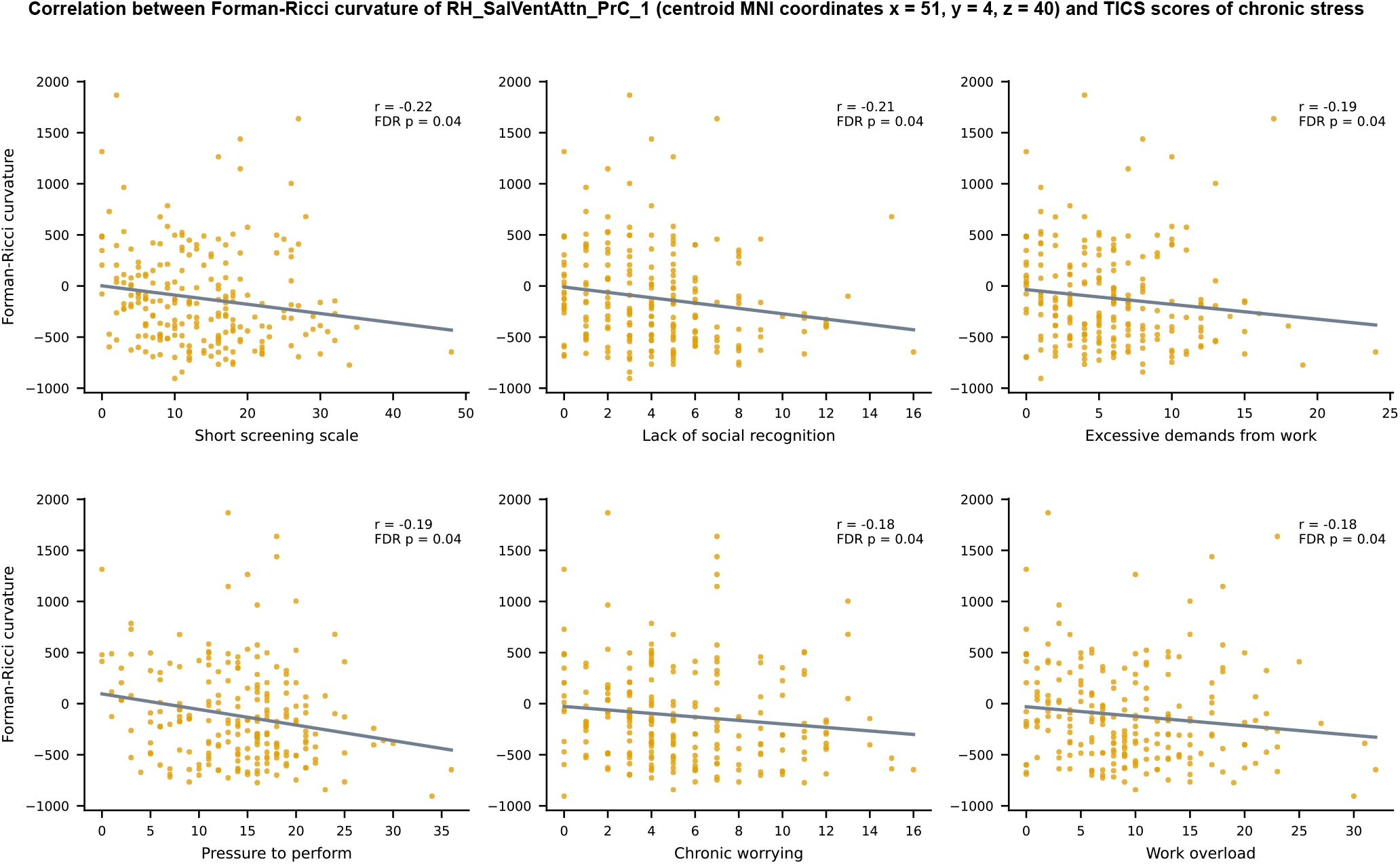
Scatter plots depicting the relationship between Forman-Ricci curvature (FRC) of the brain region RH_SalVentAttn_PrC_1 and 6 TICS scores of chronic stress namely, Short Screening Scale for Chronic Stress (SSCS), lack of social recognition, excessive demands from work, pressure to perform, chronic worrying and work overload. Note that only the scatter plots corresponding to significant correlations *r* between FRC of RH_SalVentAttn_PrC_1 and TICS scores (*p* < 0.05, FDR corrected) are shown in this figure. Each plot displays a line describing the linear relationship between FRC of RH_SalVentAttn_PrC_1 and the corresponding TICS score, estimated using the least squares method. The brain region is named according to the labelling scheme provided in the Schaefer atlas [49].

### Comparison of region-level differences in curvatures with tDCS/tACS/TMS data

The Neurosynth meta-analysis performed in this work revealed that brain regions with age-related FRC and ORC differences in somatomotor and dorsal attention networks are associated with movement. Movement is known to be impaired with age [46, 47]. Subsequently, we performed a post-hoc analysis to determine the overlap between the set of brain regions with age-related FRC and ORC differences in somatomotor and dorsal attention networks, and the set of brain regions whose stimulation with non-invasive techniques, e.g. tDCS, yielded improvement in motor function of elderly individuals. An overlap between these two sets of regions would provide additional evidence, apart from results from the Neurosynth meta-analysis, that FRC and ORC identify brain regions that are related to movement impairments in healthy elderly individuals. To perform this analysis, we first performed a literature search on PubMed to identify scientific papers reporting the effect of non-invasive brain stimulation on motor performance in healthy elderly individuals. The details of the PubMed search query are provided in the **Materials and Methods** section. **Figure 1** summarizes the workflow we employed to collect and classify the eligible articles from the literature survey, as guided by preferred reporting items for systematic reviews and meta-analysis (PRISMA) [75]. We used results reported in these eligible articles to identify those brain regions whose stimulation in elderly individuals using tDCS, tACS or TMS yielded improvement in motor performance. Finally, we compared this set of brain regions to those with age-related FRC or ORC differences in somatomotor and dorsal attention networks.

We found that previous studies applying tDCS on healthy old adults reported significant improvements in motor performance upon stimulating 4 target regions namely, primary motor cortex (M1), dorsolateral prefrontal cortex (DLPFC), right supplementary motor area (SMA) and the cerebellum. Previous studies that applied tACS on healthy old adults reported significant improvements in motor performance upon stimulating 3 target regions namely, DLPFC, posterior parietal cortex (PPC) and left M1. Finally, previous studies that applied TMS on healthy old adults reported significant improvements in motor performance upon stimulating the left DLPFC. We refer readers to **Supplementary Table S2** for details about the tDCS, tACS and TMS studies.

Note that the target regions stimulated in the previous tDCS, tACS or TMS studies correspond to ROIs in the Brodmann atlas [80, 85], whereas the ROIs in the present study are defined according to the Schaefer atlas. Thus, we mapped the ROIs in the Schaefer atlas to ROIs in the Brodmann atlas [40] (see **Materials and Methods**) to enable systematic comparison of the age-related region-level differences in discrete Ricci curvatures with those target regions whose stimulating resulted in improved motor performance. **Supplementary Table S7** lists the ROIs in the Brodmann atlas corresponding to each of the 200 ROIs in the Schaefer atlas. Across tDCS, tACS and TMS, the four cortical target regions showing evidence for improvements in motor performance in healthy elderly individuals include M1 (Brodmann area 4), DLPFC (Brodmann areas 9 and 46 [85]), PPC (Brodmann areas 5, 7, 39 and 40 [86]) and right SMA (Brodmann area 6). These target regions map to 42 ROIs in the Schaefer atlas. Out of these 42 ROIs, 11 ROIs belong to the somatomotor network and 12 ROIs belong to the dorsal attention network. All the 11 ROIs in the somatomotor network whose stimulation resulted in improved motor performance in healthy older individuals also showed age-related differences in both FRC as well as ORC. Further, 5 out of the 12 ROIs in the dorsal attention network whose stimulation resulted in improved motor performance in healthy older individuals also showed age-related differences in both FRC as well as ORC. **Figure 6** provides a visual illustration of the overlaps between the set of ROIs whose non-invasive stimulation resulted in improved motor performance in elderly, and the set of ROIs with age-related differences in FRC or ORC, within the somatomotor and dorsal attention networks. These results provide evidence that both FRC and ORC identify brain regions that are related to movement impairments in healthy elderly individuals.

**FIG. 6.**
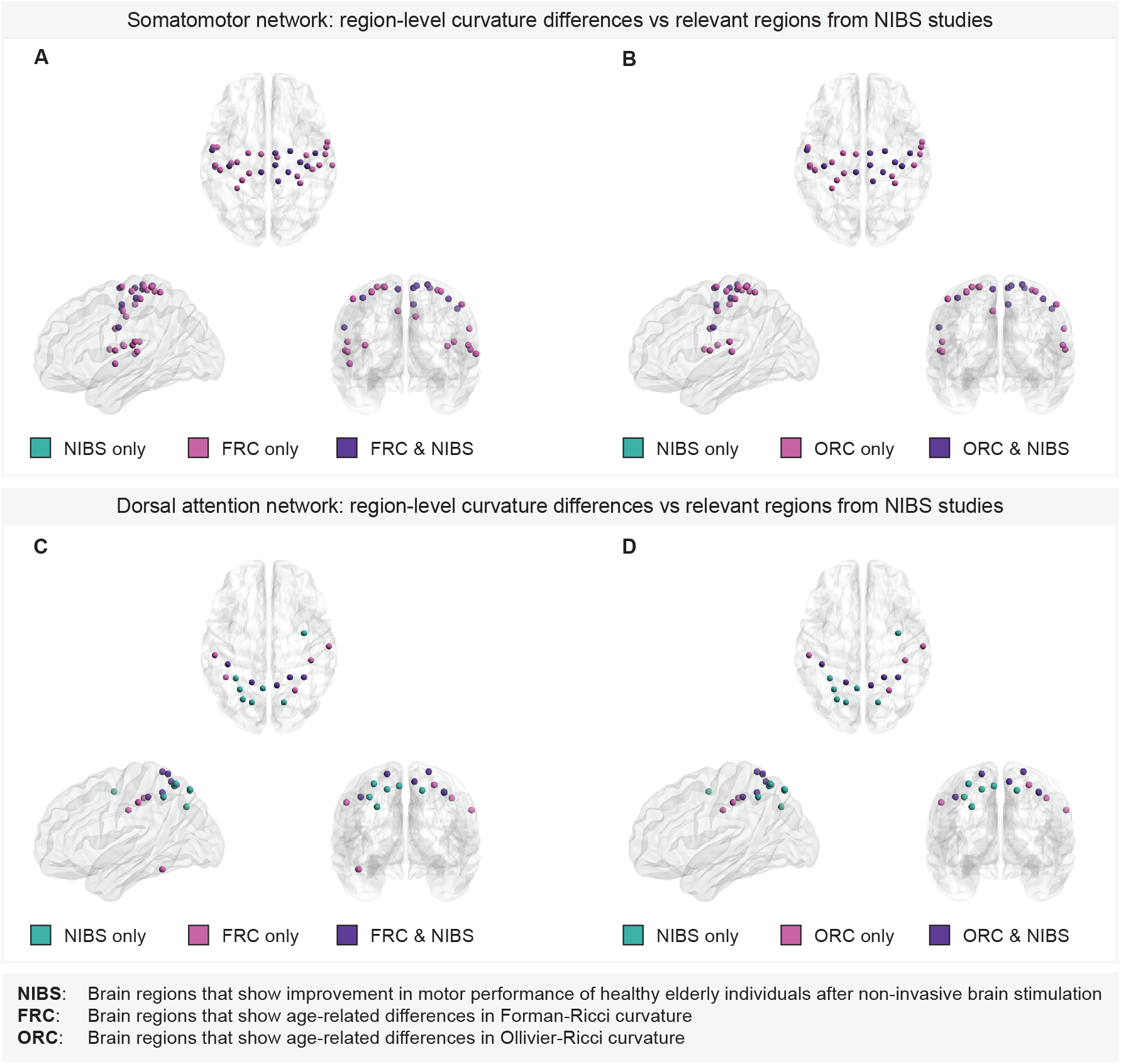
Overlap between brain regions with age-related curvature differences and target regions whose non-invasive stimulation resulted in improved motor performance of healthy elderly individuals. **(A)** Within the somatomotor network, overlap between the set of 33 Schaefer ROIs with age-related differences in Forman-Ricci curvature (FRC) and 11 Schaefer ROIs corresponding to the target regions used in tDCS/tACS/TMS experiments. All the 11 ROIs corresponding to the target regions show age-related differences in FRC. **(B)** Within the somatomotor network, overlap between the set of 27 Schaefer ROIs with age-related differences in Ollivier-Ricci curvature (ORC) and 11 Schaefer ROIs corresponding to the target regions used in tDCS/tACS/TMS experiments. All the 11 ROIs corresponding to the target regions show age-related differences in ORC. **(C)** Within the dorsal attention network, overlap between the set of 10 Schaefer ROIs with age-related differences in FRC and 12 Schaefer ROIs corresponding to the target regions used in tDCS/tACS/TMS experiments. 5 out of the 12 ROIs corresponding to the target regions show age-related differences in FRC. **(D)** Within the dorsal attention network, overlap between the set of 9 Schaefer ROIs with age-related differences in ORC and 12 Schaefer ROIs corresponding to the target regions used in tDCS/tACS/TMS experiments. 5 out of the 12 ROIs corresponding to the target regions show age-related differences in ORC. Note that the set of brain regions in each subfigure is partitioned into the following three subsets. Regions that are relevant according to non-invasive stimulation studies but do not show age-related curvature differences (labelled as “NIBS only”), regions that show age-related curvature differences but lack evidence from non-invasive stimulation studies (labelled as “FRC only” or “ORC only”), and regions that show both age-related curvature differences as well as relevance according to non-invasive stimulation studies (labelled as “FRC & NIBS” or “ORC & NIBS”). In subfigures **A** and **B**, there are no regions labelled “NIBS only” since all the regions in the somatomotor network that are relevant according to non-invasive stimulation studies also show age-related differences in FRC and ORC, respectively.

## DISCUSSION

Prior to this work, the utility of discrete Ricci curvatures in characterizing alterations in FCNs related to healthy aging remained unexplored. In the present work, we apply two widely-used notions of discrete Ricci curvature, FRC and ORC, to study resting-state functional connectivity differences in 153 healthy young and 72 healthy old subjects from the MPI-LEMON dataset [21]. Raw rs-fMRI scans for each subject were processed using a uniform preprocessing pipeline implemented using the CONN toolbox [48], and 49 FCNs at varying edge-densities between 2 – 50% were constructed for each subject. The nodes in the FCNs correspond to the 200 ROIs defined by the Schaefer atlas [49]. The rs-fMRI preprocessing pipeline and FCN construction methodology is identical to our previous work [40], wherein some of us applied discrete Ricci curvatures to compare FCNs of individuals with ASD and typically developing controls. After comparing FRC and ORC across the FCNs of young and elderly groups, we found age-related whole-brain and region-level differences in FCNs. Next, we used meta-analysis decoding to show that the brain regions with age-related differences in FRC and ORC are associated with the cognitive domains of movement, affective processing and somatosensory processing. We found that FRC of some brain regions with age-related differences exhibit significant correlation with behavioral test scores of affective processing. Finally, we showed that FRC and ORC can capture brain regions whose non-invasive stimulation is known to improve motor performance of older adults. **Figure 7** provides a summary of the methods and the main results obtained in this work.

**FIG. 7.**
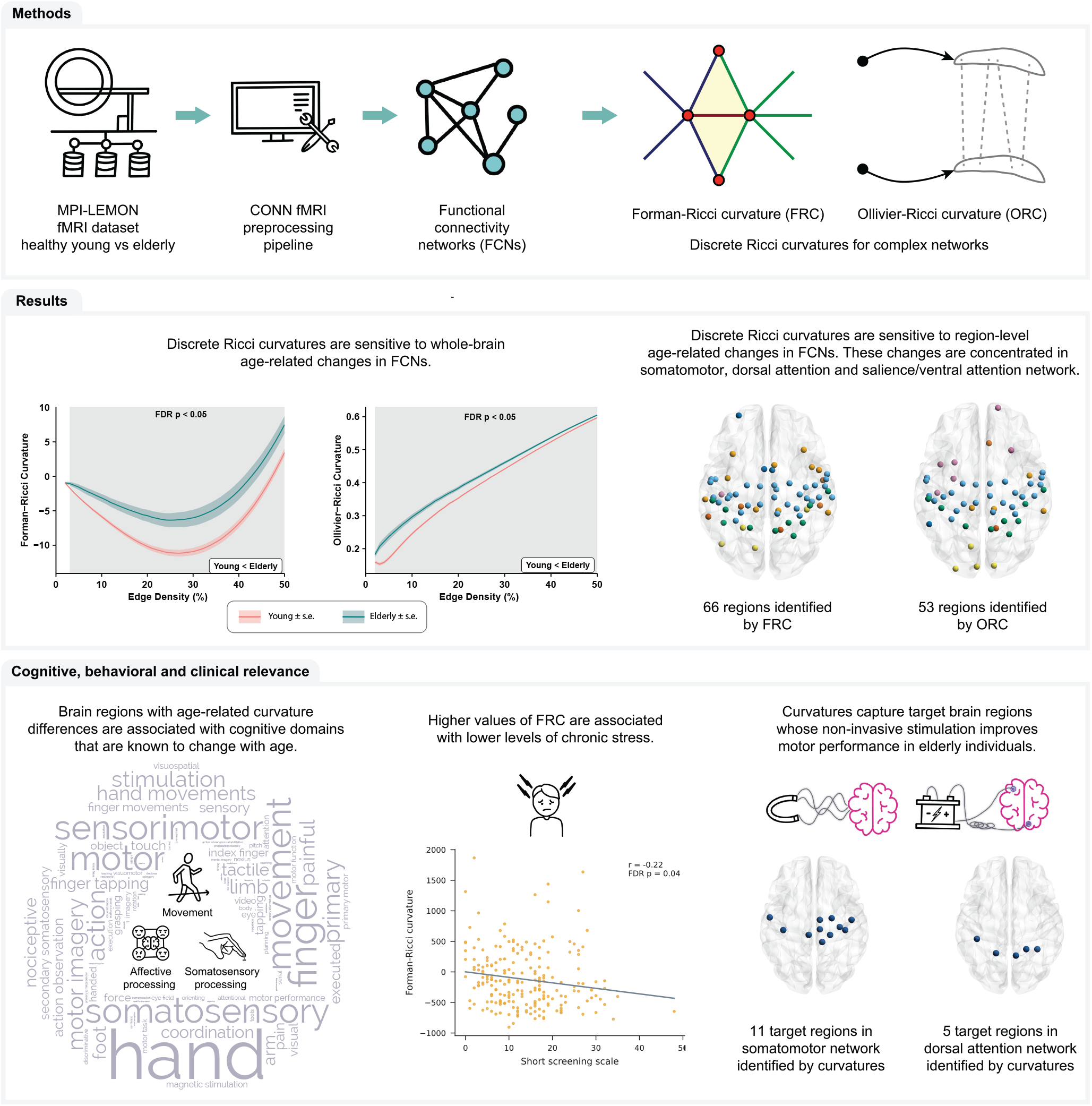
Summary of workflow and main results obtained in this study. Raw rs-fMRI scans of young participants and elderly participants were obtained from the MPI-LEMON dataset and preprocessed using the CONN toolbox. The preprocessed rs-fMRI scan of each participant was used to construct resting state functional connectivity networks (FCNs) at varying edge densities. Next, we computed and compared Forman-Ricci curvature (FRC) and Ollivier-Ricci curvature (ORC) across the FCNs of the young and elderly groups. We found that both FRC and ORC show whole-brain and region-level differences in the FCNs on young and elderly participants. Additionally, we found that age-related differences in FRC and ORC are associated with the cognitive domains of movement, affective processing and somatosensory processing. Notably, node FRC shows a significant negative correlation with behavioral test scores of chronic stress. Finally, we showed that FRC and ORC can capture brain regions whose non-invasive stimulation is known to improve motor performance of older adults.

FRC and ORC are fundamentally defined on the edges in a network. Despite being local measures by definition, the average properties of discrete Ricci curvatures have previously been used to characterize the global organization of structural and functional brain networks [37, 40] as well as financial networks [34, 35]. In this work, we compared the average edge curvatures of the FCNs in the young and elderly groups, and found that both average edge FRC and average edge ORC are significantly higher in the elderly group. Moreover, we found that the global differences in ORC exist across all the network densities considered in this work. Thus, our results suggest that discrete Ricci curvatures are highly sensitive to age-related changes in the global topological organization of resting-state FCNs.

We determined how the age-related differences in curvature are distributed across the 200 brain regions of the Schaefer atlas. We compared node FRC and node ORC across the FCNs of young and elderly individuals. We found 66 brain regions that show age-related differences in FRC and 53 brain regions that show age-related differences in ORC. Notably, FRC and ORC identify 42 brain regions in common. The age-related differences in FRC are mainly concentrated within the somatomotor, dorsal attention and salience/ventral attention network, whereas the age-related differences in ORC are concentrated within the somatomotor and dorsal attention network. Farooq *et al.* [36] have previously applied ORC to compare the structural brain networks of healthy young and healthy old individuals, and reported that brain regions with differences in ORC are present within the default mode network and visual areas. Thus, the application of discrete Ricci curvatures to functional connectivity networks can identify complementary sets of brain regions related to healthy aging, compared to those identified by applying ORC to structural brain networks.

Next, we used Neurosynth meta-analysis decoding based on a large dataset of fMRI studies to determine the cognitive and behavioral relevance of age-related differences in curvature. We found that the brain regions exhibiting age-related differences in FRC are associated with movement, affective processing and somatosensory processing, whereas the brain regions exhibiting age-related differences in ORC are associated with movement. Previous research suggests deficits in motor performance for older adults [46, 47] such as difficulties in coordination [87], increased variability in movement [88] and slowing of movement [89]. Previous research also suggests that healthy aging is associated with changes in affective processing [45]. Specifically, elderly individuals are less emotionally reactive to negative situations [45, 90] and display higher emotional maturity compared to young individuals. Further, previous literature also suggests a decline in somatosensory functions such as warmth, touch and vibration with age [91]. Notably, Lautenbacher *et al.* [92] in their systematic review and meta-analysis study have shown that pain threshold is higher in elderly individuals. Hence, our results suggest that the regions exhibiting differences in discrete Ricci curvatures in healthy aging populations are associated with the cognitive domains and abilities that are typically known to exhibit age-related changes, even in healthy aging. We remark that this is the first study that uses discrete Ricci curvatures to identify the cognitive and behavioral domains associated with age-related changes in brain networks.

After determining the cognitive and behavioral domains identified by discrete Ricci curvatures, we performed a post-hoc analysis to test the associations of discrete Ricci curvatures with altered cognition and behavior in healthy aging. We found that FRC of the nodes in the FCNs shows significant correlation with test scores related to affective processing in healthy individuals. Specifically, we found that higher values of FRC are associated with lower levels of chronic stress. These results demonstrate the ability of FRC to identify brain regions that are functionally relevant to affective processing.

Previous studies on neurocognitive aging have provided ample evidence for age-related impairments in motor performance, which may include difficulties in planning, execution and control of movement and deficits in coordination [13, 93, 94]. Non-invasive brain stimulation technologies such as tDCS, tACS and TMS provide an attractive option to modulate brain function and help preserve motor performance in older adults [76–79]. The Neurosynth meta-analysis decoding performed in the present work demonstrated the ability of both FRC and ORC to identify brain regions that are associated with movement. We performed a manual curation of non-invasive brain stimulation data to determine whether the regions with age-related differences in curvature hold any clinical significance for the treatment of motor declines in healthy elderly individuals. We found that brain regions with age-related differences in curvature overlap with those brain regions whose non-invasive stimulation with tDCS, tACS or TMS shows evidence for improvement in the motor performance of healthy elderly. Remarkably, within the somatomotor network, both FRC and ORC were able to detect the brain regions that are clinically relevant according to non-invasive stimulation experiments. These results suggest that discrete Ricci curvatures can be used to generate novel hypothesis about target regions for non-invasive brain stimulation experiments.

We highlight a methodological aspect of applying discrete Ricci curvatures to brain networks and place our results in a broader context. In this work, we found similar results for FRC and ORC while determining brain regions related to healthy aging. However, it is important to note that the two notions differ in their definitions as they capture different properties of the classical Ricci curvature, and while both notions are shown to be correlated for many empirical networks [27], one may perform better than the other depending on the type of network under consideration. For example, we previously applied FRC and ORC to identify brain regions related to atypical resting state functional connectivity in autism spectrum disorder [40]. We found that FRC is able to identify more regions, provides better interpretability in terms of behavior, and can detect clinically significant regions that are not captured by ORC. Therefore, future works on the application of discrete Ricci curvatures to brain networks may benefit when different notions of curvature are used together, as it may help gather complementary information about the topological organization of brain networks. Notably, in the present study, we found that FRC can capture more brain regions related to healthy aging compared to ORC within the salience/ventral attention network, whereas ORC is able to capture more regions in the limbic network. Further, at the level of cognitive domains, we found that FRC identifies brain regions related to three domains, namely movement, affective processing and somatosensory processing, whereas ORC identifies brain regions related only to movement.

To summarize, we found that geometry-inspired notions of discrete Ricci curvature can be used to characterize age-related changes in brain functional connectivity, both at the whole-brain level as well as at the level of individual brain regions. Further, we showed that the brain regions captured by curvatures hold clinical relevance for non-invasive brain stimulation interventions that are focused towards preserving motor function of older adults. Future studies could expand beyond healthy aging populations by applying discrete Ricci curvatures to brain networks of age-related neurodegenerative disorders such as Alzheimer’s disease and Parkinson’s disease, and derive insights about the cognitive domains affected in age-related neurodegenerative disorders.

## Acknowledgements

We would like to thank Kishan Kumar for help with figures. A.S. would like to acknowledge research support from the Max Planck Society, Germany via a Max Planck Partner Group in Mathematical Biology and the Department of Atomic Energy (DAE), Government of India. J.J. acknowledges support from the German-Israeli Foundation (GIF) grant number I-1514-304.6/2019.

## Author contributions

A.S., Y.Y., P.E. and N.W. designed research. Y.Y. and P.E. performed the computations. Y.Y., P.E., N.W., J.J. and A.S. analyzed data and wrote the paper. Y.Y. and P.E. generated the figures. A.S. supervised the project.

## Ethics statement

The human participants included in the MPI-LEMON dataset provided written informed consent prior to any data acquisition, including agreement to share their data anonymously. Further, the original MPI-LEMON study was carried out in accordance with the Declaration of Helsinki and the study protocol was approved by the ethics committee at the medical faculty of the University of Leipzig (Reference number 154/13-ff).

## Data and Code Availability

Functional connectivity matrices and networks generated in this study, and all original computer programs are available via the GitHub repository: https://github.com/asamallab/Curvature-FCN-Aging.

