## Supplementary Figure for "Discrete Ricci curvatures capture age-related changes in human brain functional connectivity networks"

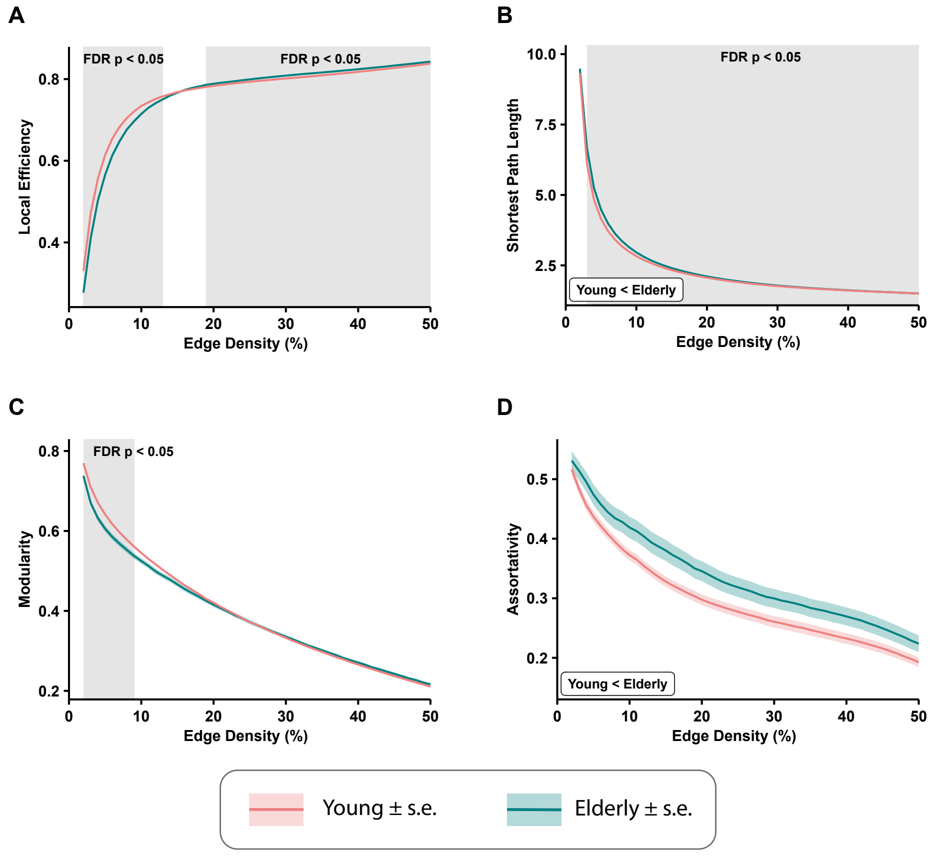


FIG. S1. Differences in standard global network measures across the functional connectivity networks (FCNs) of 153 young individuals and 72 elderly individuals. The differences are reported for FCNs over the range of edge densities 0.02 (i.e., 2% edges) and 0.5 (i.e., 50% edges), with an increment of 0.01 (i.e., 1% edges). The shaded regions in each plot correspond to the edge densities where the between-group differences are statistically significant ($p < 0.05$, FDR-corrected). **(A)** Average local efficiency is significantly higher in young individuals over edge densities $1 - 12\%$, and significantly higher in elderly individuals over edge densities $19 - 49\%$. **(B)** Average shortest path length is significantly higher in elderly individuals over edge densities $3 - 49\%$. **(C)** Modularity is significantly higher in young individuals over edge densities $2 - 9\%$. **(D)** Assortativity is higher in the young group, but the differences are not statistically significant.

---

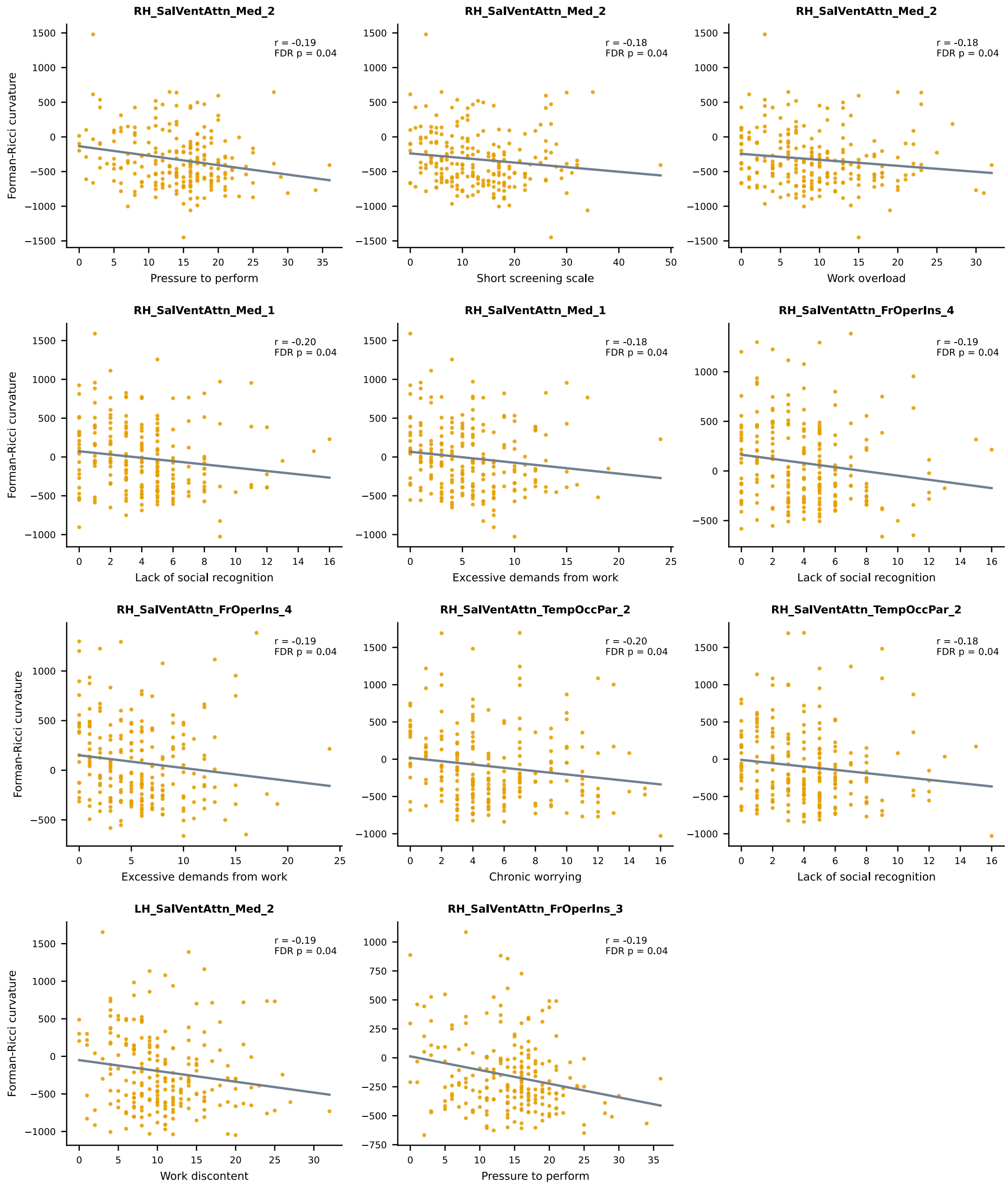


FIG. S2. Correlation between Forman-Ricci curvature (FRC) and TICS scores of chronic stress. Scatter plots depicting the relationship between FRC of 6 brain regions (RH_SalVentAttn_Med_2, RH_SalVentAttn_Med_1, RH_SalVentAttn_FrOperIns_4, RH_SalVentAttn_TempOccPar_2, LH_SalVentAttn_Med_2 and LH_SalVentAttn_FrOperIns_3), and TICS scores of chronic stress. Note that only the scatter plots corresponding to significant correlations $r$ between FRC and TICS scores ($p < 0.05$, FDR corrected) are shown in this figure. Each plot displays a line describing the linear relationship between FRC of a brain region and the corresponding TICS score, estimated using the least squares method. Brain regions are named according to the labelling scheme provided in the Schaefer atlas.
