## Supplementary Text for "Discrete Ricci curvatures capture age-related changes in human brain functional connectivity networks"

### STANDARD NETWORK MEASURES

All the standard network measures are defined for an unweighted and undirected graph $G = (V, E)$. The graph $G$ can also be denoted as $n \times n$ adjacency matrix $\mathbf{A}$, where $A_{ij} = 1$ if nodes $i$ and $j$ are adjacent, and $A_{ij} = 0$ otherwise.

- The *clique number* of $G$ is defined as the size of the maximal clique appearing in $G$.
- For a node $i$ in the graph $G$, *clustering coefficient* is defined as:

  $$C_i = \frac{2}{k_i(k_i - 1)} \sum_{j,k} (A_{ij} A_{ik} A_{jk})^{1/3}$$

  where $j$ and $k$ are the neighbors of node $i$ and the summation is over all pairs of neighbors of $i$. The *average clustering coefficent* of $G$ is the average of the clustering coefficients of the individual nodes in $G$.
- The *shortest path length* $d(i, j)$ between any two nodes $i$ and $j$ in $G$ is equal to the number of edges contained in the shortest path connecting them. *Average shortest path length* of $G$ is the average of the shortest path length between all pairs of nodes in $G$, i.e.,

  $$\langle L \rangle = \frac{1}{n(n-1)} \sum_{i,j \in V} d(i, j)$$

- *Global efficiency* measures the ability of the network $G$ to exchange information [1], and is defined as

  $$E_{glob}(G) = \frac{1}{n(n-1)} \sum_{i \neq j \in V} \frac{1}{d(i, j)}.$$

- Let $G_i$ denote the subgraph of the neighbors of node $i$ in $G$. The *local efficiency* [1] of node $i$, $E_{loc}(G_i)$ is defined as the efficiency of the subgraph $G_i$. *Average local efficiency* of $G$ is given by

  $$E_{loc}(G) = \frac{1}{n} \sum_{i \in V} E_{loc}(G_i)$$

- The *betweenness centrality* of a node $i$ in $G$ measures the extent to which it lies on the shortest path between other nodes, and is defined as [2]

  $$C_b(i) = \sum_{j,k \in V} \frac{\sigma(j, k|i)}{\sigma(j, k)},$$

  where $\sigma(j, k)$ is the number of shortest paths between $j$ and $k$, and $\sigma(j, k|i)$ is the number of shortest paths between $j$ and $k$ that pass through $i$.

- Many networks display a tendency to exhibit a modular structure, where the set of nodes in a network can be partitioned into subsets of densely connected nodes. *Modularity* of the graph $G$ measures the density of intra-module edges compared to inter-module edges, and is defined as [3, 4]

$$Q = \frac{1}{2m_w} \sum_{i \neq j \in V} [A_{ij} - \frac{s_i s_j}{2m_w}] \delta(c_i, c_j)$$

  where $s_i$ and $s_j$ give the sum of weights of edges attached to nodes $i$ and $j$, respectively, $c_i$ and $c_j$ are the communities of $i$ and $j$, respectively, and $m_w$ is the sum of all edge weights. Since $G$ is an unweighted graph, all edges in $G$ are assigned weight equal to 1.

- A network is said to be assortative when two nodes attached to a given edge in the network tend to have the same degree. The *assortativity coefficient* [5] measures the degree correlations between nodes in an unweighted network $G$, and is defined as

$$r = \frac{\sum_i e_{ii} - \sum_i a_i b_i}{1 - \sum_i a_i b_i}.$$

  Here, $e_{ij}$ is the fraction of edges in $G$ that are attached to nodes of degree $i$ and $j$. Note that $\sum_{ij} e_{ij} = 1$. Further, $a_i$ and $b_i$ are the fraction of edges whose one end is attached to nodes of degree $i$, that is $a_i = \sum_j e_{ij}$ and $b_i = \sum_j e_{ji}$. Since $G$ is an undirected network, we have $e_{ij} = e_{ji}$ and $a_i = b_i$.
